# Targeting of the nuclear RNA exosome to chromatin by HP1 affects the transcriptional programs of liver cells

**DOI:** 10.1101/2025.02.14.638307

**Authors:** Hiba Souaifan, Mickael Costallat, Laura Sitkiewicz, Kylian Godest, Florence Cammas, Carl Mann, Christian Muchardt, Christophe Rachez

**Affiliations:** Institut de Biologie Paris-Seine (IBPS), Sorbonne Université, CNRS UMR7238, 75252 Paris, France; Institute of Human Genetics, CNRS UMR9002 University of Montpellier, 34396 Montpellier, France; Institut de Biologie Intégrative de la Cellule (I2BC), CEA, CNRS, Université Paris-Saclay, 91190 Gif-sur-Yvette, France; Ecole doctorale Complexité du Vivant, Sorbonne Université, Paris, France

## Abstract

HP1, a hallmark of pericentromeric heterochromatin, is a chromatin-bound regulator of co-transcriptional processes including alternative splicing, but its role in RNA degradation remains unexplored. Here, we uncover a direct interaction between HP1 and the RNA exosome, a major RNA decay complex. In mouse embryonic liver cells, inactivation of all three HP1 isoforms led to accumulation of retrotransposon-derived RNAs and stabilization of enhancer RNAs. These changes coincided with increased activity at a subset of liver enhancers particularly sensitive to reduced exosome activity, many of which regulate genes encoding extracellular matrix components such as *Col6a1* and *Col6a2*. Stratifying hepatocellular carcinoma samples by HP1 expression further revealed that tumors with low HP1 were marked by reduced RNA degradation, and increased expression of a similar subset of genes encoding extracellular matrix components and possibly contributing to tumor stiffness. These results suggest that HP1’s impact on RNA turnover contributes to its function in cancer biology.

## Introduction

Chromatin structure is a major regulator of transcription. This regulation is mediated by molecular machineries that establish transcriptionally active or repressed chromatin states, known as euchromatin and heterochromatin, respectively. In active regions, chromatin influences gene expression programs that govern processes ranging from development to responses to external stimuli. In contrast, constitutive heterochromatin is typically found at pericentromeric regions and transposable elements, and is enriched with repressive epigenetic marks— notably di- and tri-methylation of histone H3 on lysine 9 (H3K9me2/3). These marks serve as binding sites for HP1 proteins, which are essential for the establishment, maintenance, and propagation of heterochromatin ^1^.

The three mammalian HP1 isoforms—HP1α, HP1β, and HP1γ—share a conserved structure consisting of two globular domains separated by an intrinsically disordered hinge region. The N-terminal chromodomain specifically recognizes and binds H3K9me2/3-marked nucleosomes, anchoring HP1 to heterochromatic regions ^2–4^. The C-terminal chromoshadow domain mediates dimerization and interactions with a broad range of PxVxL motif–containing proteins involved in transcriptional regulation, DNA replication, and repair ^5, 6^. The unstructured hinge region has an affinity for nucleic acids ^7^. In addition to their well-established role in heterochromatin, HP1 proteins also localize to actively transcribed genes in both *Drosophila* and mammals, with isoform-specific patterns ^8–10^. In euchromatin, HP1 proteins have been implicated in several co-transcriptional processes, including the regulation of alternative splicing ^11–15^, recruitment of RNAi machinery to chromatin ^12^, and control of transcriptional elongation ^16^ and termination ^17^.

Multiple lines of evidence indicate that the RNA-binding activity of HP1 proteins underlies their functions in both euchromatin and heterochromatin. In both compartments, HP1 proteins engage with repeated RNA sequences. This was first demonstrated by the requirement of RNA binding for HP1 targeting to heterochromatin ^18, 19^. Later, it was further shown that HP1α specifically associates with pericentromeric major satellite repeat RNA (SatRNA) ^20^. More recently, we showed that HP1γ contributes to splicing precision by recognizing hexameric motifs enriched in intronic SINE elements, supporting intron–exon discrimination and contributing to accurate splice site definition ^21, 22^. In *Schizosaccharomyces pombe*, recognition of repeated sequences by the HP1 homolog Swi6 has been further linked to the degradation of satellite SatRNAs, contributing to transcriptional repression beyond chromatin condensation. Two distinct mechanisms may mediate this repression. First, Swi6 is required for RNAi-dependent silencing by directing SatRNA to the RITS complex, enabling siRNA production via double-stranded RNA formation ^23^. Second, silencing at centromeric repeats and the silent mating-type locus—both heterochromatic regions—also involves polyadenylation-dependent RNA degradation via the RNA exosome cofactor Cid14 ^24^. Swi6 then engages in a dynamic exchange between free, chromatin-bound, and RNA-bound states, facilitating the delivery of repeat-derived RNAs to degradation machineries ^25^.

The RNA exosome complex is a central machinery for RNA degradation and quality control in eukaryotic cells ^26^. Its core structure consists of nine non-catalytic subunits (Exosc1–9) that form a scaffold for associated 3′→5′ ribonucleases. In the cytoplasm, the exosome partners with the ribonuclease Dis3l and cytoplasmic cofactors to form the SKI complex, which mediates mRNA turnover and surveillance ^27^. In the nucleus, the exosome associates with two ribonucleases: Rrp6/Exosc10, a distributive exonuclease, and Rrp44/Dis3, a processive exo- and endonuclease. The nuclear exosome functions within specialized complexes—NEXT, PAXT, and TRAMP—that provide substrate specificity across a broad spectrum of nuclear RNAs ^28^. The TRAMP complex, well characterized in *S. cerevisiae*, contributes to ribosomal RNA processing and snoRNA precursor degradation. In mammals, however, TRAMP is primarily restricted to the nucleolus ^29^. By contrast, the NEXT complex targets unprocessed RNAs with a free 3′ end, such as promoter-upstream transcripts (PROMPTs), uaRNAs, and enhancer RNAs (eRNAs) ^30–32^, while the PAXT complex directs degradation of polyadenylated nuclear RNAs ^33^. A defining feature of all exosome complexes is the presence of an RNA helicase. In the cytoplasm, this activity is provided by Ski2, while in the nucleus, Mtr4 is essential for substrate recruitment to NEXT, PAXT, and TRAMP (reviewed in ^34^. Recruitment of these nuclear complexes is further mediated by the cap-binding complex (CBC), which recognizes the 5′ methyl-guanosine cap of RNA polymerase II transcripts. CBC, along with its cofactor ARS2, connects to NEXT via the zinc-finger protein ZC3H18, forming the CBC–NEXT axis ^35^. ZC3H18 bridges the 5′ cap to the exosome, facilitating the degradation of unstable or aberrant RNAs.

The activity of the RNA exosome has previously been linked to the organization of chromatin structure. At a large scale, chromatin architecture, particularly topologically associating domains (TADs) and chromatin loops, is shaped by architectural proteins such as CTCF and cohesins ^36^. Depletion of the nuclear exosome subunit DIS3 impairs non-coding RNA processing, reduces CTCF binding, and disrupts TAD organization at the *Igh* locus ^37^. In parallel, transcripts derived from normally silenced retroelements are targeted for degradation by the NEXT complex, which recognizes unstable RNAs arising from heterochromatic regions ^38, 39^. Mass spectrometry (MS) approaches have finally suggested that RNA exosome-associated factors such as ZC3H18 and the RNA helicase MTR4 may associate with HP1 ^40, 41^, while, reciprocally, HP1 MS data has detected ZC3H18 among the interacting partners of all three HP1 isoforms ^42^. This association suggest that molecular bridges may exist between chromatin, HP1, and nuclear RNA degradation machineries.

Building on this possible connection, we have here analyzed the relationship between chromatin-associated HP1 proteins and RNA exosome components in murine BMEL liver cell lines depleted of all three HP1 isoforms. Previous studies using these cells showed that HP1 depletion results in the accumulation of transcripts from normally silenced retroelements, including LINEs, SINEs, and LTR-containing endogenous retroviruses (ERVs) ^43, 44^. In this context, we confirm physical associations between HP1 proteins and several representative components of the nuclear exosome, and show an HP1-dependent enrichment of exosome complexes at chromatin loci. HP1 loss also led to a marked increase in chromatin-associated unstable transcripts, such as PROMPTs and retroelement-derived RNAs, particularly at LTR loci. Notably, some of these transcripts extended beyond the annotated repeat boundaries and were detected in a mature form in the cytoplasm. In parallel, enhancers located near these retroelements became activated in the absence of HP1, leading to upregulation of adjacent genes and widespread alterations in transcriptional programs.

## Results

### HP1 proteins interact with components of the RNA Exosome and affect their subcellular localization

Analysis of ZC3H18 immunoprecipitation–mass spectrometry data ^41^ using the STRING database (Search Tool for the Retrieval of Interacting Genes/Proteins) documented that the protein-protein interactions linking the RNA exosome and the Cap binding complex with the three HP1 proteins and other epigenetic regulators, were supported by multiple independent sources (Fig. 1A, each “string” represents a documented protein-protein interaction). This prompted us to further investigate the links between HP1 proteins and the nuclear RNA Exosome.

**Fig. 1:**
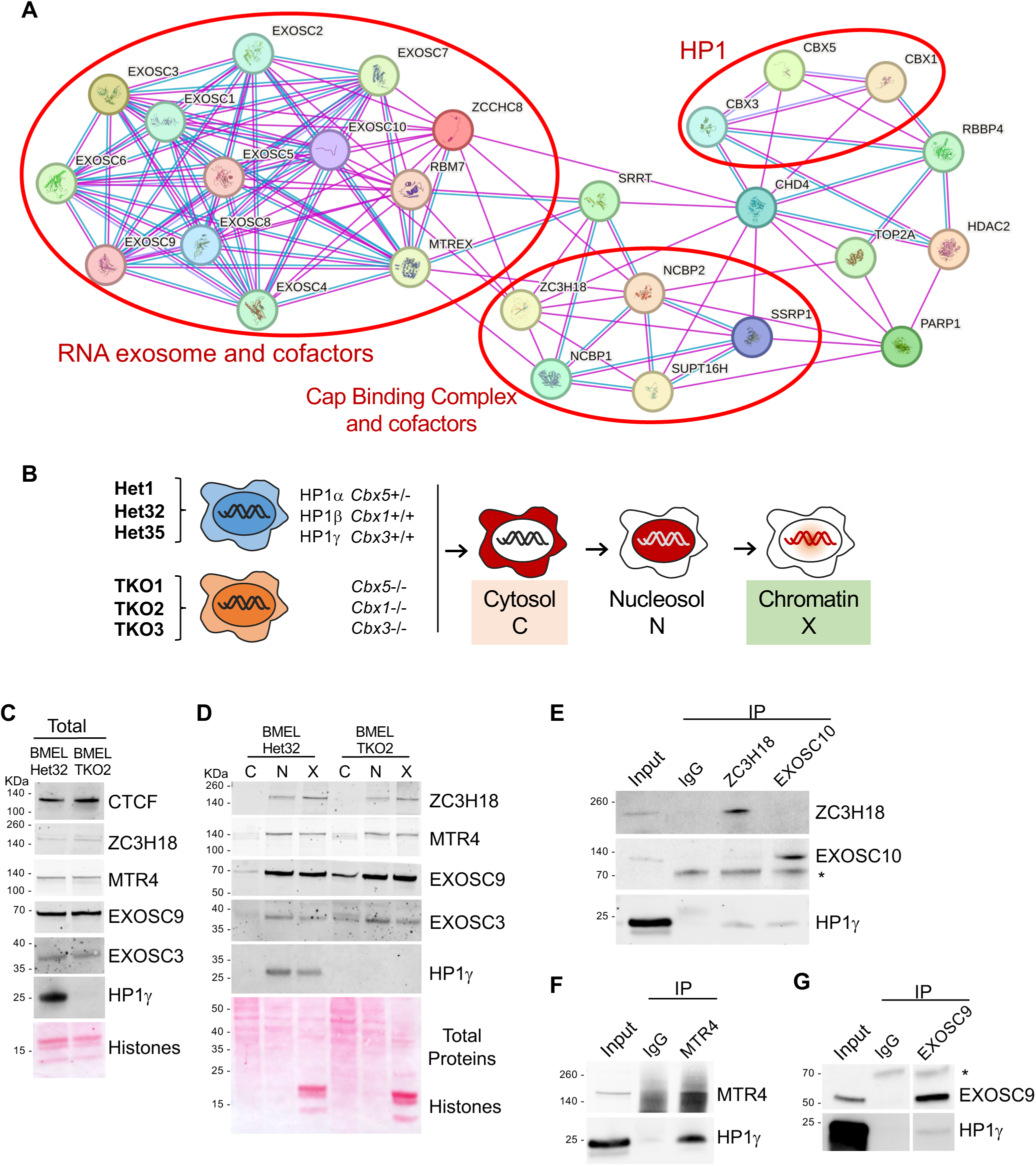
Protein-protein association between HP1 and the RNA Exosome. A. String analysis of known physical proteins associations among factors co-precipitating with ZC3H18 proteins, identified by interaction profiling of ZC3H18, in ref. ^41^. Pink and blue network edges correspond to protein associations based on experiments or curated databases, respectively. **B.** Genotype and subcellular fractionation scheme of the six BMEL clones expressing (Het) or not (Triple Knock-out, TKO) HP1 proteins used in this study. **C.** Protein levels in total extracts, and **D,** subcellular distribution in the three fractions, cytosolic (C), nucleosoluble (N), and chromatin (X), depicted in Figure 1B of indicated exosome subunits and cofactors in Het and TKO cells by Western blot analysis and Ponceau staining of total proteins. **E, F, G.** Co-immunoprecipitation of HP1γ by MTR4, ZC3H18, EXOSC10, or EXOSC9 in BMEL nuclear extracts.

To this end, we compared mouse embryonic liver (BMEL) cells lacking all three HP1 isoforms (triple knockout, TKO) to HP1-expressing controls (Het, heterozygous for HP1α and wild type for HP1β and HP1γ). Six independent clones were analyzed: TKO1, TKO2, TKO3, and Het1, Het32, Het35 (Fig. 1B and Supplementary Fig. 1A). Western blot analysis of total extracts showed that protein levels of EXOSC3 and EXOSC9 (core exosome subunits), MTR4 (a nuclear RNA helicase common to all nuclear exosome complexes), and ZC3H18 (a zinc-finger adaptor protein specific to the NEXT complex) were similar between Het and TKO clones, indicating that HP1 depletion did not affect their overall expression (see representative clones, Fig. 1C). Upon fractionation of the Het clones into cytosolic (C), nucleosolic (N), and chromatin (X) extracts, the four proteins were predominantly detected in the nucleosolic and chromatin fractions, which were also enriched in HP1γ (Fig. 1D). Interestingly, in the TKO clones, we observed increased accumulation of EXOSC9 and EXOSC3 in the cytosolic fraction, suggesting reduced nuclear retention of these subunits (Fig. 1D, lanes C). Fractionation quality was validated by Western blotting for marker proteins and by analyzing the distribution of representative cytoplasmic and nuclear transcripts (Supplementary Fig. 1A–C).

Finally, co-immunoprecipitation from nuclear extracts containing both soluble and chromatin-associated material confirmed an association of HP1 proteins with EXOSC9, EXOSC10, MTR4, and ZC3H18, consistent with the previously reported MS data (Fig. 1E-G and Supplementary Fig. 1D-E). These interactions were maintained following RNase treatment (Supplementary Fig. 1F), indicating that they were not mediated by RNA. Together, these results further documented interactions between the HP1 proteins and components of the RNA exosome machinery, and suggested that these interactions are required for full nuclear retention of certain exosome subunits.

### Several non-coding, unstable RNA species are found stabilized in the absence of HP1 proteins, in both chromatin and cytosolic fractions

We next used RNA-seq to examine RNA profiles in chromatin and cytosolic fractions of the BMEL Het and TKO clones (Fig. 1B). In this setup, the chromatin fraction was expected to be enriched in nascent and incompletely processed RNAs, whereas the cytosolic fraction was expected to be depleted of unstable nuclear transcripts and enriched in mature mRNAs (Supplementary Fig. 1C). In parallel, RNA-seq was also performed on total RNA from each clone. The RNA species we examined included various RNAs encoded by LTRs, LINEs, SINEs, and MMSAT repeats. For these, we focused on extragenic repeat elements, excluding those located within gene bodies, such as intronic repeats, which may be transcribed incidentally as part of coding gene expression and processed in that context. We also examined levels of upstream antisense uaRNA at promoters (PROMPT RNA), and eRNA at active distal enhancers (enhD RNA), all documented as unstable transcripts. Genomic elements were considered as expressed when yielding at least two RNA-seq reads in at least one experimental condition.

Using this criterion, levels of unstable transcripts increased in TKO cells compared to HP1-expressing (Het) cells. In particular, we noted that for more than 50% of all expressed LTR elements, expression was detected exclusively in TKO cells (see examples in Fig. 2A, and quantification in Fig. 2B). Several elements were also expressed exclusively in Het cells, reflecting the multiplicity of HP1 functions not solely linked to RNA stability ^45^. Global analysis of RNA abundance further showed that mean RNA levels from PROMPTs, distal enhancers (enhD), and repeat elements (LTRs, LINEs, SINEs) were increased in the chromatin fraction of TKO cells relative to Het (Fig. 2C). In contrast, chromatin-associated pre-mRNAs from Ensembl-annotated genes, measured by exonic reads, were less affected by HP1 depletion. Globally, the Ensembl gene transcript category showed more moderate changes in both RNA levels and the number of loci with TKO- or Het-specific expression compared to non-coding RNA categories (Fig. 2E-G). These findings suggest that HP1 loss predominantly affects the expression and/or stability of unstable, non-coding RNAs, rather than processed mRNAs. Supporting this, several non-coding RNAs that accumulated in the chromatin fraction—particularly those derived from LTRs, LINEs, and MMSAT repeats—were also found at higher levels in the cytoplasm of TKO cells (Fig. 2D). In addition, many LTR-initiated transcripts extended for several kilobases and, interestingly, some appeared to be spliced in the cytoplasm (e.g., RLTR17, RMER13A; Fig. 2A), suggesting that in the absence of HP1, these RNAs evade nuclear decay and undergo canonical maturation.

**Fig. 2:**
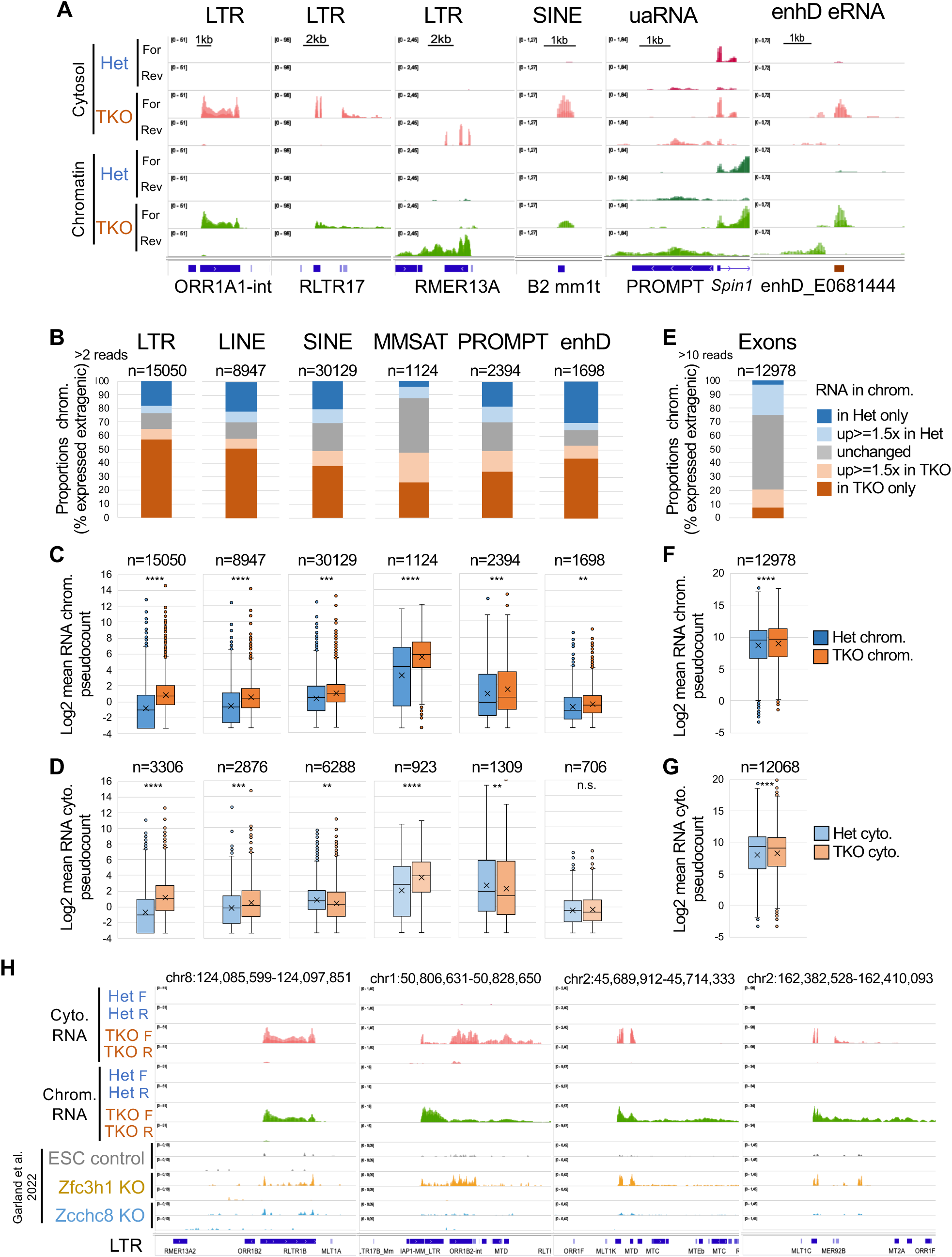
Chromatin-enriched RNA is increased in TKO cells and stabilized in the cytoplasm. **A.** Genome views of RNA density profiles (merged triplicates) in chromatin (green) and cytosolic (red) fractions, at indicated genomic loci showing TKO-specific or TKO-increased RNA. **B, C, D.** RNA levels by counting uniquely mapped reads on repeated elements (LTR, LINE, SINE, MMSAT4), Promoter upstream antisense RNA (PROMPT), distal enhancers (enhD) and exons of ensembl genes. **B.** Cumulative histograms of proportions of the indicated genomic regions expressing RNA (more than 2 reads) by comparison of the three Het clones with the three TKO clones. Five categories discriminate expression on chromatin in TKO only (brown), increased in TKO (salmon), in Het only (dark blue), increased in Het (light blue), unchanged (grey). **C, D.** Boxplots of mean RNA read counts in chromatin (C) or in the cytosol (D) on genomic regions as in B. in Het (blue) or TKO (orange). Asterisks represent significance level by unpaired two-tailed Student’s t-test (**** p < 0.0001, *** p < 0.001, ** p < 0.01, * p < 0.05, n.s., not significant). **E, F, G.** Histogram and boxplots as in figures B, C, D, for proportions and counts on expressed exons (more than 10 reads). **H.** Transcriptome profiles in Het or TKO chromatin (green) or cytoplasm (red) compared with transcriptomes of mESC invalidated for the expression of NEXT or PAXT Exosome complex components, *Zcchc8* KO (sky blue profiles) or *Zfc3h1* KO (yellow profiles) and control mESC (grey profiles) (GSE178550; Garland et al., 2022). All boxplots in Fig 2 show median (central line), IQR (25th and 75th percentiles, box limits); whiskers extend to 1.5 x IQR.; points are values of outliers.

Notably, several of the LTR-derived transcripts most readily detected in TKO cells (Fig. 2H, green and red profiles) were also reported as overexpressed in a previous study following depletion of ZFC3H1 or ZCCHC8, components of the PAXT and NEXT nuclear exosome pathways, respectively, in mouse embryonic stem cells ^38^; Fig. 2H, yellow and light blue profiles). While derived from a different cell type, this overlap supports the idea that the accumulation of LTR transcripts observed in TKO cells reflects impaired exosome-mediated RNA degradation. Together, these results suggest that HP1 loss stabilizes normally unstable non-coding RNAs, possibly due to defective degradation at the chromatin level.

### An HP1-dependent association of nuclear RNA exosome and cofactors to chromatin loci

Having observed that HP1 inactivation alters the nuclear retention of RNA exosome components and leads to the accumulation of unstable RNAs on chromatin, we next sought to examine more closely how RNA exosome components associate with nuclear structures. We therefore performed chromatin immunoprecipitation followed by high-throughput sequencing (ChIP-seq) with anti-EXOSC10, -ZC3H18, and -MTR4 antibodies using the Het32 and the TKO2 clones. To also gain insight in the overall impact of HP1 inactivation on chromatin structure, we completed this series of experiments with anti-CTCF, and anti-phospho RNA polymerase II (P-RNAPII, as a mix of both phospho-Ser2 and -Ser5 forms) ChIP-seq, and HP1α, HP1ϕ3, HP1γ, H3K9me3, and H3K27me3 CUT&Tag experiments (Fig. 3A).

**Fig. 3:**
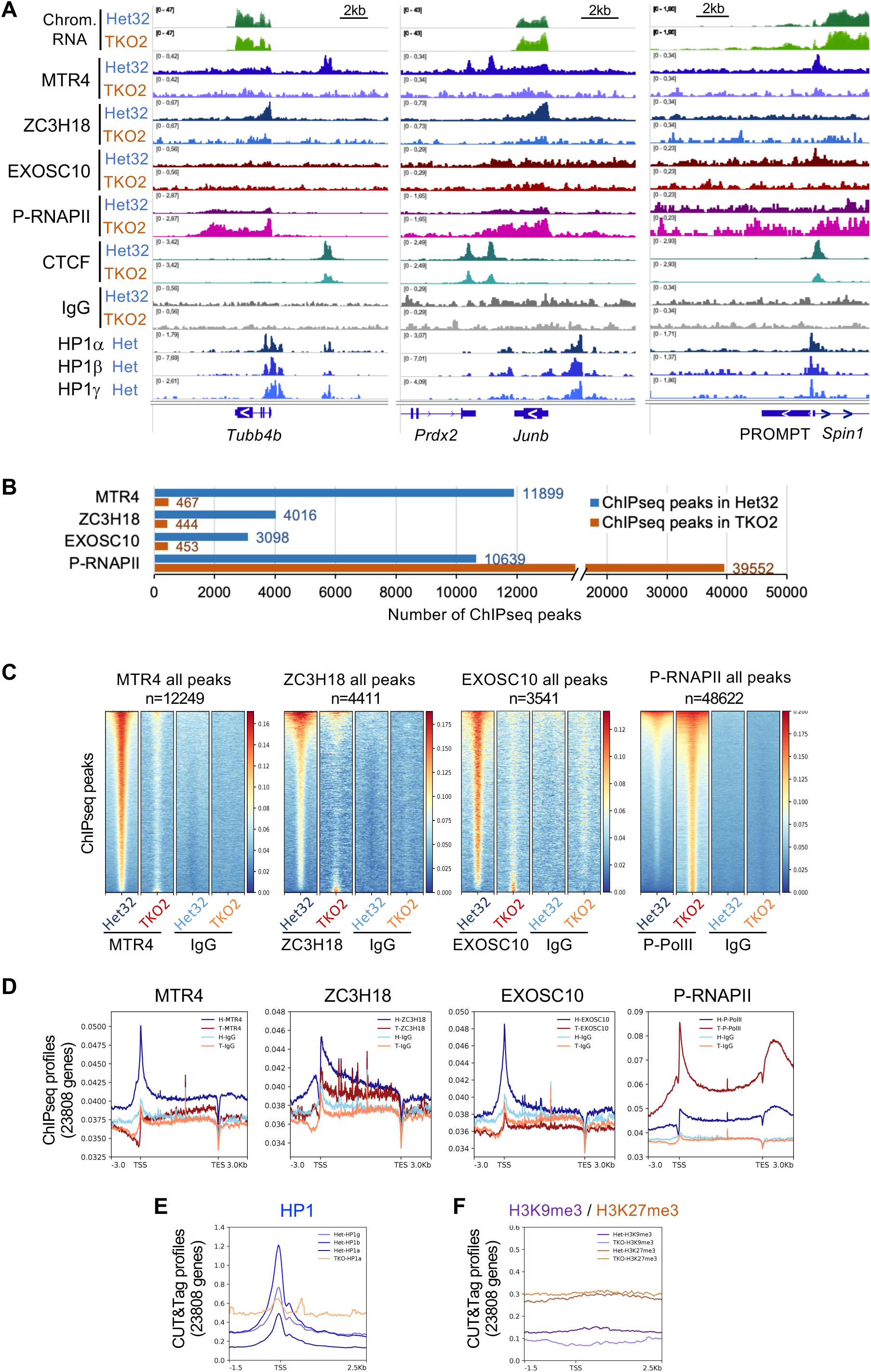
ChIPseq analysis reveals an HP1-dependent targeting of Nuclear RNA exosome and cofactors to chromatin loci. **A.** Genome views of chromatin RNA density profiles (top), ChIPseq profiles of indicated proteins, and HP1 CUT&Tag density profiles (bottom) on indicated loci. **B.** Number of ChIPseq peaks versus IgG defined by MACS2 analysis for the indicated proteins in Het32 (blue) and TKO2 (brown). **C.** Heatmaps of genome-wide ChIPseq densities centered on ChIPseq peaks in both Het32 and TKO2 combined in a single list (all peaks). **D.** Metagene analysis of ChIPseq profiles of the indicated proteins on 23808 gene bodies defined as TSS to TES intervals, in Het32 (dark blue), in TKO2 (dark red), compared to IgG profiles in Het32 and TKO2 (salmon and light blue profiles, respectively). **E.** CUT&Tag profiles at gene TSS for the three HP1 proteins (E) in Het (blue profiles), and HP1a in TKO (salmon profile). **F.** Cut&Tag profiles at TSS for H3K9me3 (purple) and H3K27me3 (brown) histone marks in Het (dark colors) and in TKO (light colors).

Examination of the ChIP-seq profiles with a genome browser revealed that the nuclear exosome components, MTR4, ZC3H18, and EXOSC10 are actively recruited to chromatin at specific genomic loci. This recruitment was most clearly detected in regions also recruiting P-RNAPII, while Mtr4 also displayed an interesting overlap with CTCF. Importantly, the chromatin recruitment of exosome components was strongly diminished in TKO2 cells, indicating that HP1 proteins are required for their proper localization to these regions. In contrast, levels of phosphorylated RNA polymerase II (P-RNAPII) appeared increased in TKO2 cells (Fig. 3A). Quantification of MTR4, ZC3H18, and EXOSC10 ChIP-seq peaks detected with the MACS2 package confirmed a dramatic decrease in the number and/or intensity of these peaks upon inactivation of the HP1 genes, while the total number of P-RNAPII peaks was strongly increased (Fig. 3B-C).

Metagene analysis on more than twenty thousand genes confirmed a preferential enrichment of MTR4, ZC3H18, EXOSC10 on the body of genes, with a peak at the transcription start site (TSS - Fig. 3D). Thus, the distribution of these factors coincided with that of the P-RNAPII, although missing accumulation at sites of transcription termination (TES). This analysis further revealed that HP1 inactivation led to a uniform reduction in RNA exosome component-occupancy over gene bodies, while RNAPII occupancy was correspondingly increased (Fig. 3D, compare brown and dark blue profiles). CUT&Tag analysis of the recruitment of each HP1 isoform in Het cells showed their preferential enrichment at gene transcriptional start sites (TSS, blue profiles Fig. 3E), compared to the background HP1 alpha CUT&Tag signal in TKO cells (orange profile Fig. 3E). This distribution of HP1 proteins at actively transcribed genes is consistent with earlier reports^46^. It is also compatible with a co-recruitment of HP1 and RNA exosome proteins at promoter regions. Thus, HP1 inactivation leads to both reduced recruitment of the RNA degradation machinery and increased recruitment of RNA polymerase II, supporting a model in which HP1 proteins contribute to gene silencing by simultaneously restricting transcription and promoting RNA degradation. Of note, although HP1 enrichment at TSS was not associated with H3K9me3 or H3K27me3 marks, HP1 colocalized with H3K9me3 at other regions, such as simple repeats (Fig. 3E and 3F, and Supplementary Fig. 3A and 3B). At these sites, HP1 inactivation did not lead to a loss of H3K9me3 but rather to a redistribution of the mark across repeats. These observations suggest that the targeting of the RNA exosome to TSS by HP1 and its association with H3K9me3 at other loci rely on two independent mechanisms.

### HP1 concentrates MTR4 and CTCF at gene promoters

We next examined in more detail the apparent colocalization of MTR4 with CTCF peaks. Quantification confirmed that 90% of the MTR4 peaks match CTCF peaks (representing approximately 15% of the total number of CTCF peaks). It also showed that MTR4 was the only RNA exosome component under scrutiny to display this colocalization (Fig. 4A). Comparison of ChIP-seq data between Het32 and TKO2 cells showed that the overall number of CTCF peaks remained broadly unchanged following HP1 depletion (Fig. 4B). However, genome browser visualization revealed a redistribution of CTCF binding: in TKO2 cells, new CTCF peaks appeared at previously annotated loci where no binding was detected in Het32 cells, while many pre-existing peaks decreased in intensity, eventually below levels of detection (Fig. 4C). Heatmap analyses confirmed this genome-wide redistribution of CTCF occupancy (Fig. 4D). The genome-browser visualization further suggested that a large fraction of the CTCF peaks colocalizing with MTR4 were located at promoters. Sorting promoters according to HP1 occupancy revealed that CTCF and MTR4 were co-enriched at HP1-bound promoters in Het32 cells and that HP1 inactivation led to a strong reduction of both CTCF and MTR4 enrichment at these sites, accompanied by an increase in P-RNAPII signal (Fig. 4E). Together, these results suggest that HP1 proteins contribute to concentrating both CTCF and MTR4 at a subset of promoters, possibly also preventing it from ectopic associating with other CTCF binding sites.

**Fig. 4:**
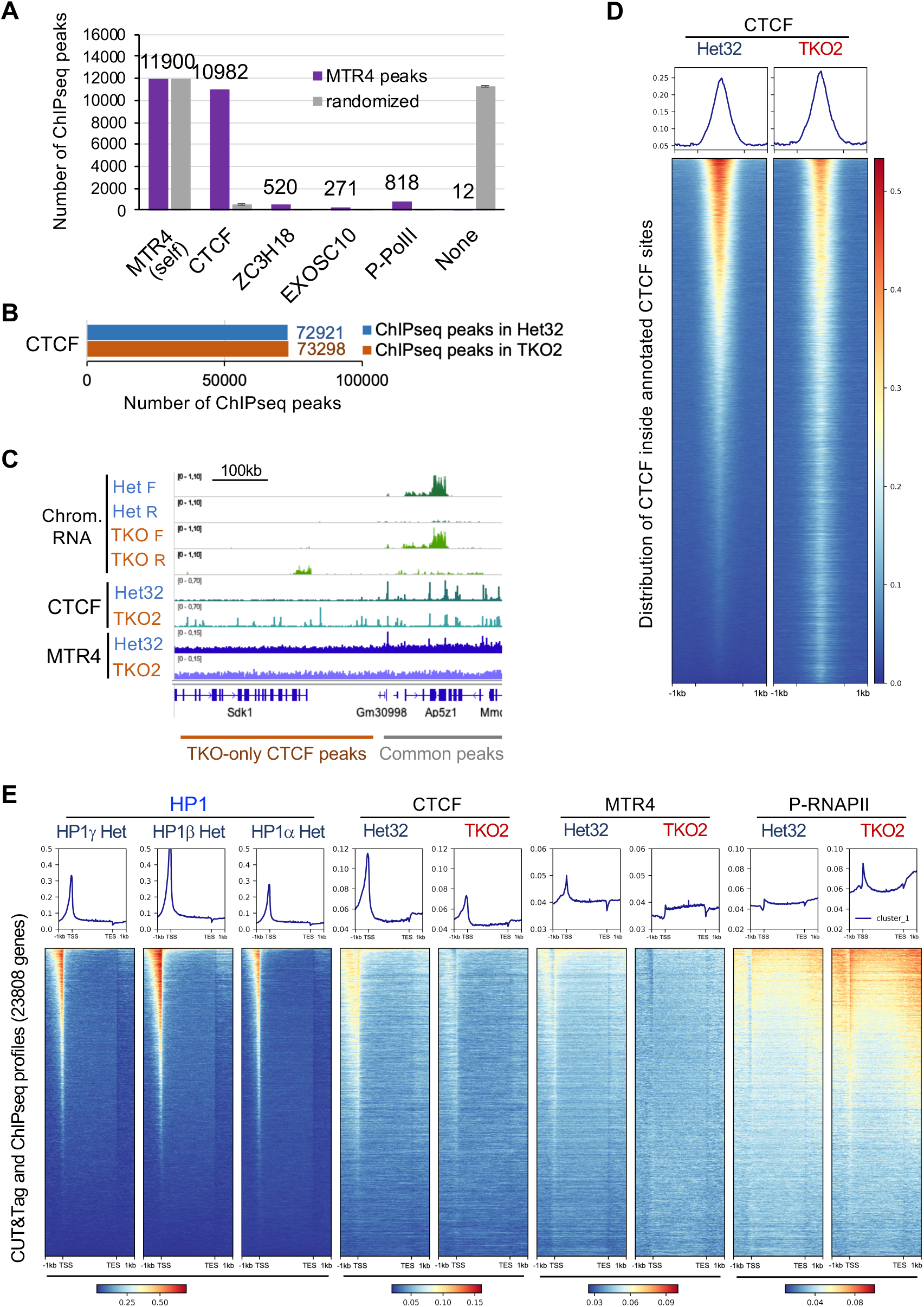
HP1-dependent CTCF and MTR4 at promoters. **A.** Number of MTR4 peaks (purple bars) colocalized with the other indicated ChIPseq peaks, compared to the same list of MTR4 peaks with randomized genomic locations (gray bars). **B,** Number of CTCF ChIPseq peaks versus IgG defined by MACS2 analysis in Het32 (blue) and TKO2 (brown). **C.** Genome view of chromatin-enriched RNA, CTCF and MTR4 densities at a locus showing two regions bearing TKO-only CTCF peaks (on the left), or peaks common to both cell types (on the right). **D.** Heatmaps of genome-wide CTCF ChIPseq densities in Het32 and TKO2 centered on all CTCF ChIPseq peaks. **E.** Heatmaps of ChIPseq densities of the indicated proteins on 23808 gene bodies defined as TSS to TES intervals, in Het32 (dark blue), in TKO2 (dark red).

### A gain in chromatin accessibility at enhancers affects collagen gene expression

To gain broader insight into the regulatory elements impacted by HP1 inactivation, we intersected both CTCF ChIP-seq peaks and HP1 CUT&Tag peaks with the catalog of candidate cis-regulatory elements (cCREs) annotated by the ENCODE project (Fig. 5A and Supplementary Fig. 5A). This analysis revealed that CTCF peaks were located within annotated regulatory elements, including promoters and enhancers, both locations where they were mostly lost in the absence of HP1 (Supplementary Fig. 5A). In parallel, approximately 40% of HP1 peaks were also located within promoters as well as enhancers, with a predominance of distal (enhD) versus promoter-proximal enhancers (enhP) (Fig. 5A). To further investigate the consequences of HP1 loss on enhancer activity in BMEL cells, we performed ATAC-seq in Het and TKO clones. Based on chromatin accessibility profiles, distal enhancers (enhD) were categorized as active in both conditions, with decreased activity in TKO cells, with increased activity in TKO cells, or inactive (Unchanged, Down, Up, Inactive, respectively. Supplementary Fig. 5B). Heatmap analyses revealed that, similar to observations at promoters, a substantial fraction of active enhancers were associated with HP1 in Het cells (Fig. 5B).

**Fig. 5:**
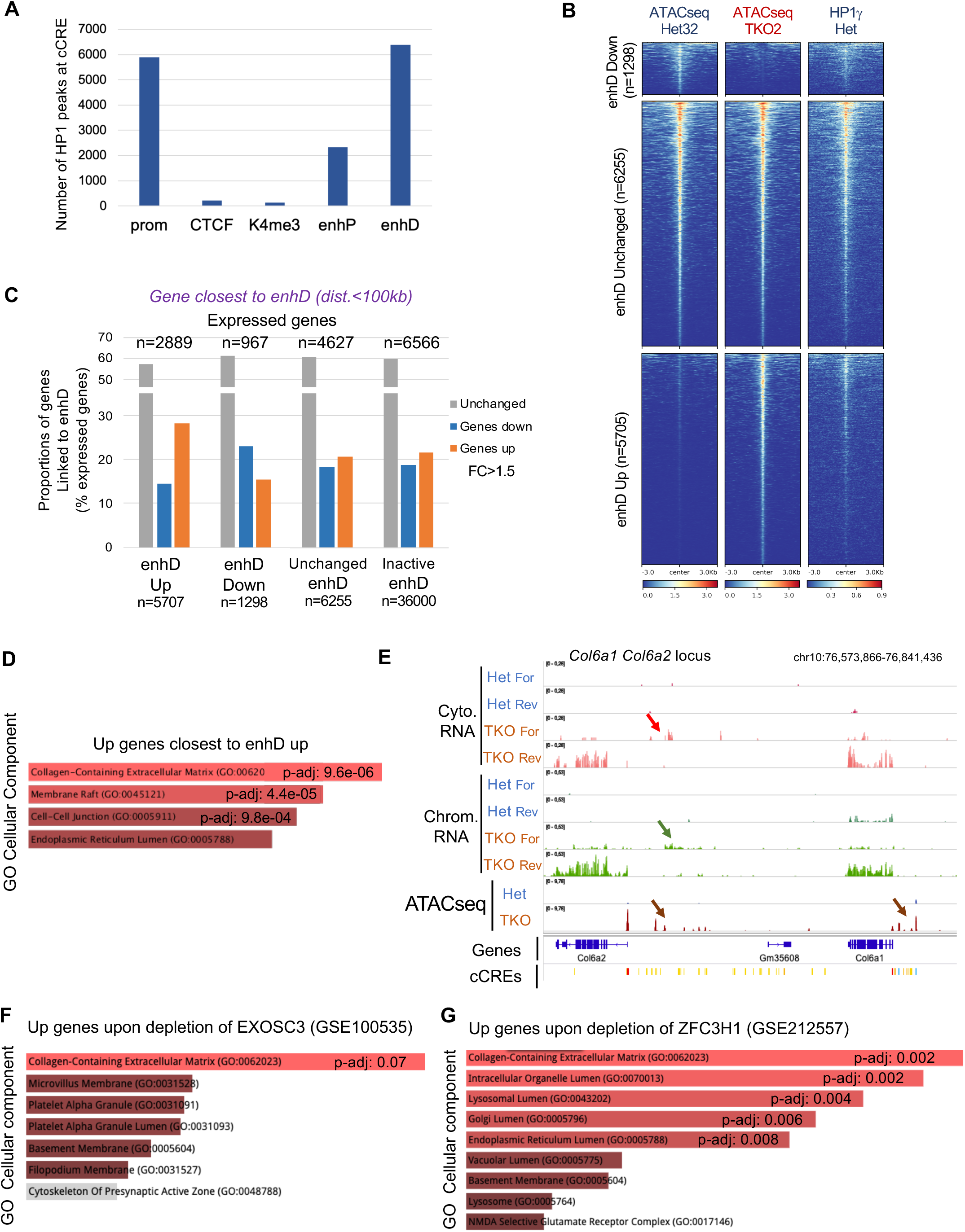
Changes in chromatin accessibility at enhancers affect collagen gene expression. **A.** Number of CUT&Tag HP1 merged peaks located at cCRE regulatory elements annotated as in Supp. Fig. 5A. **B.** Heatmaps of ATACseq densities in Het and TKO cells, and of HP1g CUT&Tag density in Het cells centered at cCRE enhD enhancer regulatory elements. EnhD were categorized based on their chromatin accessibility by ATACseq between Het and TKO. Categories are as follows: Down, with decreased accessibility in TKO cells; Unchanged, active in both conditions; Up, with increased accessibility in TKO cells; inactive (not shown), with no ATACseq density, respectively. **C.** Proportions of expressed genes in the closest proximity (within 100kb) to enhD in the categories depicted in 5B. Bars represent genes that were up-(orange) or down-regulated (blue) by more than 1.5 fold in TKO versus Het, or unchanged (grey). The total number of genes associated with each enhD category is indicated above the histogram. **D.** GO term analysis by enrichR associated with upregulated genes closest to upregulated enhD. **E.** Genome view of stranded chromatin-enriched (green) and cytosolic (red) RNA densities on the *Col6a1 Col6a2* locus, together with ATACseq densities in Het and TKO, showing TKO-specific ATACseq density (brown arrows) colocalizing with TKO-specific RNA (green and red arrows) and a group of enhD elements depicted in the cCRE lane. Yellow bars, enhD/enhP; red bars, Prom; blue bar, CTCF, as annotated in Supp.Fig. 5A. **F, G.** GO term analysis by enrichR associated with upregulated genes upon depletion of the indicated RNA exosome components.

Importantly, activation of enhancers upon HP1 inactivation correlated with the upregulation of nearby genes located within 100 kb (enhD Up, orange bar, Fig. 5C). Gene ontology analysis of these upregulated genes revealed a strong enrichment for terms associated with the collagen-containing extracellular matrix (Fig. 5D). Further examination showed that multiple collagen isoforms were specifically upregulated at enhancers derepressed upon HP1 inactivation, as exemplified at the *Col6* locus (Fig. 5E, and list in Supplementary Fig. 5C). This activation correlated with an increased accumulation of transcripts over the enhD elements in the chromatin fraction, and at some loci, also in the cytosolic fraction, suggesting increased stability of enhancer RNAs (eRNAs)—potentially a consequence of the reduced recruitment of RNA exosome components to chromatin (Fig. 5E, arrows; Sup. Fig. 5B). This mechanism may also be at play for the few thousand LTR retroelement transcripts that were stabilized in TKO (Sup. Fig. 5D). These findings suggest that collagen genes are particularly sensitive to decreased RNA exosome activity. To further support this hypothesis, we analyzed publicly available transcriptomic datasets from *Exosc3* conditional knockout mouse embryonic stem cells (GSE100535, Fig. 5F) and HeLa cells depleted of ZFC3H1 (GSE212557, Fig. 5G). Remarkably, in both models, genes associated with the collagen-containing extracellular matrix were among the most significantly affected pathways following disruption of RNA exosome activity.

### HP1 Deficiency and Reduced RNA Exosome Activity Converge to Deregulate Collagen Genes in Hepatocellular Carcinoma

To investigate the relevance of HP1–RNA exosome cooperation in hepatocellular carcinoma (HCC), we reanalyzed RNA-sequencing data from a cohort of 76 Mongolian HCC patients ^47^. The molecular subtypes identified in this cohort largely mirrored those previously described in patients from other geographic regions, including Asia, Europe, and North America. Our analysis revealed that HP1 gene expression—including CBX3, CBX1, and CBX5—was upregulated in the majority of tumors, with 71% of cases showing at least a twofold increase relative to adjacent healthy tissues (Supplementary Fig. 6A). This observation is consistent with previous reports linking elevated HP1 levels to poor prognosis in HCC (Supplementary Fig. 6B).

To examine the consequences of differential HP1 expression, we ranked patients based on cumulative HP1 levels (sum of *CBX3*, *CBX1*, and *CBX5* expression) and stratified them into Hi HP1 and Lo HP1 groups, corresponding to the upper and lower quartiles (Fig. 6A; Supplementary Fig. 6C). This stratification effectively separated the cohort into two subgroups in unsupervised PCA (Fig. 6B). The Hi HP1 group was enriched for patients previously associated with poor prognosis and included a higher proportion of deceased individuals (Fig. 6A; Supplementary Fig. 6D–E; ref. 47). Genes upregulated in Hi HP1 tumors were significantly enriched for cell cycle– related pathways (adjusted p-value = 4.87e-14; Supplementary Fig. 6F). These tumors also showed increased expression of the alpha-fetoprotein gene *AFP* and *YAP1*, markers of poorly differentiated and highly proliferative hepatocytes (Fig. 6D, Lo HP1 vs Hi HP1). Together, these data suggest that Hi HP1 tumors are more aggressive, with enhanced proliferation and loss of hepatocytic identity.

**Fig. 6:**
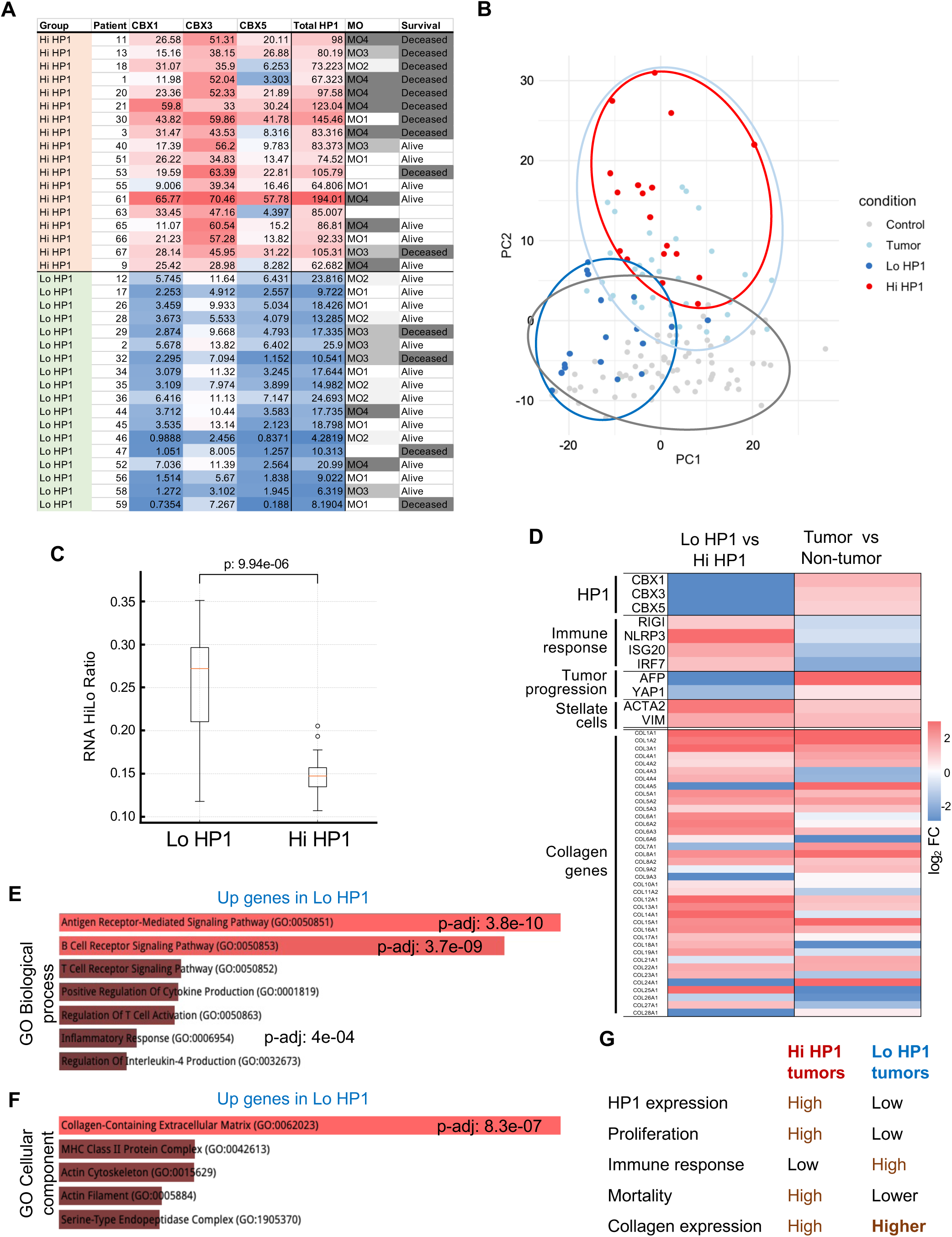
HP1 expression affects collagen genes in hepatocellular carcinoma (HCC) patient samples. **A.** Table of patient ranking in high HP1 (Hi HP1) and Low HP1 (Lo HP1) expressing samples based on their cumulative total HP1 expression levels, and compared with gene expression of individual HP1 isoforms. Molecular subclasses (MO) sorted by increasing severity of the disease from MO1 to MO4, and patient survival criteria have been analyzed in Candia *et al.*, 2020. **B.** Unsupervised PCA analysis of patient samples. **C.** Quantification of the accumulation of unstable mRNAs in Lo HP1 and Hi HP1 groups using the RNA HiLo ratio analysis as in ref. ^48^. ***, p-value : 9.94E-06 (two-tailed Student’s t-test). **D.** Heatmaps of log2 fold changes in expression of genes representative of tumor properties indicated as categories on the left and described in the main text, in Lo HP1 versus Hi HP1 tumor samples (left) and in tumor versus matching non-tumor samples (right). Log2 FC were obtained by DESeq2 analysis. **E, F.** Histograms of the most significant indicated Gene Ontology (GO) terms and their adjusted p-values for upregulated genes in Lo HP1 patient samples. **G.** Summary of tumor characteristics in the Hi and Lo HP1 groups.

To estimate RNA turnover in the Lo HP1 group, we quantified the accumulation of unstable mRNAs using the RNA HiLo ratio—a validated proxy for RNA degradation activity ^48^. This approach was used because the available transcriptomes lacked sufficient depth to detect rare non-coding RNAs from enhancers or promoters directly. The analysis revealed that labile mRNA species accumulated significantly more in Lo HP1 tumors compared to Hi HP1 tumors, consistent with reduced RNA degradation (Fig. 6C). This accumulation correlated with increased expression of antiviral response genes, including *RIG-I*, *NLRP3*, *ISG20*, and *IRF7* (Fig. 6D, Lo HP1 vs Hi HP1), suggesting activation of an RNA-triggered innate immune response. Supporting this, we observed elevated expression of genes predominantly expressed in immune cells rather than hepatocytes, indicating that the innate immune activation is accompanied by inflammation in the tumor microenvironment.

In parallel, GO cellular component analysis identified “collagen-containing extracellular matrix” as the top enriched category (Fig. 6F). This is consistent with our findings in mouse liver cells with reduced RNA exosome activity (Fig. 5D). In Lo HP1 tumors, we also observed increased expression of α-SMA/*ACTA2* and vimentin, markers of activated hepatic stellate cells (HSCs) (Fig. 6D). These changes suggest an increased fibrogenic state, in which HSCs secrete extracellular matrix proteins and drive fibrotic remodeling of the tumor microenvironment.

An increase in extracellular matrix (ECM) stiffness is a characteristics of tumoral progression of HCC, where increased ECM is preventing immune infiltration in the tumor, allowing HCC to reach more severe stages ^49^. Indeed tumor samples compared to their matching non-tumor controls revealed typical high levels of collagen gene expression and low immune response (Fig 6D, tumor vs. non-tumor). However our findings reveal that Lo HP1 tumors promote both a fibrotic evolution and an anti-viral innate immune response affecting the tumor microenvironment (Summarized in Fig. 6G). We state that this immune response is likely to be triggered by the accumulation of undegraded RNA and is key to the better survival of Lo HP1 HCC patients. They further imply that the increased expression of collagen-associated genes may be a direct consequence of reduced HP1/RNA exosome activity on chromatin, and that these genes are particularly sensitive to enhancer regulation dependent on chromatin-linked RNA degradation mechanisms.

## Discussion

We identified an unrecognized role for chromatin-associated HP1 proteins in guiding the nuclear RNA exosome to specific genomic loci. HP1 depletion in BMEL cells led to the accumulation of normally unstable nuclear RNAs, including transcripts from repetitive elements, promoters, and enhancers. The increased production of eRNAs was also accompanied by the activation of a subset of enhancers and the subsequent upregulation of nearby genes, many of which were involved in the extracellular matrix, including several collagen genes. This had important functional consequences in the context of hepatocellular carcinoma.

The RNA profile of HP1-depleted cells recapitulated the effects observed upon depletion of nuclear RNA exosome components. PROMPTs and eRNAs were stabilized both in our HP1-deficient cells and in cells lacking core exosome (EXOSC3), NEXT (ZCCHC8), or PAXT (ZFC3H1) subunits, as reported in previous studies ^50^ ^33^. In addition, ChIP-seq analysis revealed a dramatic reduction in the number of exosome-binding peaks across the genome in TKO cells, indicating a failure in local recruitment. This suggested that HP1 cooperated with NEXT and/or PAXT to promote exosome-mediated RNA decay by directing exosome components to specific chromatin loci. In this context, we noted that HP1 loss did not displace the RNA exosome from chromatin globally, nor did it affect RNA exosome subunit expression, supporting a direct role in targeting rather than chromatin association or complex stability.

The LTR-derived transcripts specifically detected in HP1 triple knockout (TKO) cells point to a dysregulation of RNA exosome activity. LTRs, derived from transposable elements (TEs), are typically embedded in heterochromatin and remain transcriptionally silent in most mammalian cells. This silencing is maintained through several mechanisms: DNA methylation predominates in somatic tissues and late embryonic stages, while in embryonic stem cells, H3K9me3-dependent pathways play a major role, particularly in repressing younger, more transcriptionally active retroelements ^51^. Parallels can be drawn from other systems. In *S. pombe*, heterochromatic silencing involves two HP1 homologs: Chp2 represses transcription via the SHREC histone deacetylase complex, while Swi6 binds centromeric noncoding RNAs and promotes their degradation via the RNAi pathway or the RNA exosome ^23, 25^. In mESC, the human silencing hub (HUSH) complex contributes to TE silencing by being recruited to H3K9me3-marked chromatin through its MPP8 subunit. HUSH also cooperates with RNA degradation pathways, including the NEXT complex, to repress TEs ^38, 52^. Although we cannot fully exclude a loss of transcriptional repression in HP1-depleted BMEL cells, our data suggest that impaired RNA degradation is the dominant factor. Specifically, the LTR loci showing increased transcript accumulation in TKO cells did not overlap with HUSH-regulated loci identified by MPP8 ChIP-seq in mESCs (Supplementary Fig. 3C-D), making HUSH involvement in this context unlikely. However, other silencing factors, such as TNRC18—an H3K9me3 reader implicated in silencing of young ERV loci ^53^ — could still contribute. Overall, the increased LTR-derived RNA levels observed in TKO cells likely reflect a combined effect: the loss of HP1-mediated transcriptional repression and defective RNA degradation. This dual mechanism would resemble what has been described in yeast (SHREC and RNAi/exosome) and mESCs (HUSH and NEXT). This hypothesis awaits further investigation.

Although retroelements like LTRs have lost their ability to transpose in mammalian genomes, they still retain many transcriptional regulatory elements due to their viral origin. These elements can serve as platforms for transcription factor binding and initiation, thereby influencing various gene regulatory mechanisms ^54^. Over time, many LTRs have been co-opted as alternative promoters, enhancers, or even chromatin insulators, such as at CTCF binding sites. In our previous work, we identified several examples of LTR exaptation as enhancers in BMEL cells ^43^, and we have also shown that increased transcription of ERVs in multiple sclerosis may reflect reactivation of embryonic enhancers of retroviral origin ^55^. In the current study, we found that a relatively small subset of LTR elements gave rise to stabilized transcripts in HP1-deficient (TKO) cells. This suggests that HP1 proteins do not broadly repress LTRs, but instead act on transcriptionally active LTRs, particularly those with enhancer potential, much like the RNAi machinery, which targets loci based on RNA production. In our data, RNAs accumulating from enhancers and transcriptionally active LTRs were associated with increased transcription of nearby genes. While it remains debated whether eRNAs act through their transcription or as functional RNA molecules, they have been proposed to promote chromatin accessibility and modulate the recruitment of transcription factors and elongation regulators ^56^. Their accumulation in HP1-depleted cells may thus contribute to the observed increase in RNAPII recruitment/phosphorylation, possibly by sequestering the negative elongation factor NELF and promoting the activation of P-TEFb ^57^.

Interestingly, genes involved in collagen and extracellular matrix formation were particularly sensitive to reduced RNA exosome activity. Both HP1 depletion and impaired exosome function were associated with their upregulation, and these genes also showed increased activity in HCC patients with low HP1 expression. The basis for this specificity remains unclear, but certain eRNAs and long noncoding RNAs (lncRNAs) are especially prone to exosome-mediated degradation and accumulate when the exosome is inactivated ^58^. It is possible that enhancers regulating extracellular matrix genes belong to this susceptible class. From a disease perspective, overexpression of both HP1 and EXOSC components has been associated with poor prognosis in cancers such as HCC (Supplementary Fig. 6B and ref. ^59^). Paradoxically, complete HP1 loss promotes liver tumorigenesis in aged mice ^44^. We hypothesize that HP1 depletion leads to RNA accumulation, triggering antiviral immune responses. While this response may offer short-term protection and reduce initial tumor severity, its chronic activation could promote inflammation and tumor progression over time, particularly in the aging liver.

## Methods

### Cell lines

Het and TKO cells (Het 1, Het 3.2, Het 3.5, TKO1, TKO2, TKO3), as described in Figure 1A, were independent clones of BMEL cells derived from mouse embryonic liver, generated as previously described ^44^. The cells were cultured in RPMI 1640 medium (Invitrogen) supplemented with 10% fetal bovine serum (FBS), 1% penicillin-streptomycin, 10 µg/mL insulin, 30 ng/mL IGF-II, and 50 ng/mL EGF, at 37 °C under 5% CO2.

### Cell fractionation and total cell extracts

Cells (approximately 10⁶ from a confluent 100 mm plate) were washed twice with ice-cold PBS directly on the plate then incubated on ice for 8 minutes in 10 mL of swelling buffer (SW): 10 mM Tris-HCl pH7.5, 2 mM MgCl2, 3 mM CaCl2, supplemented before use with 1x antiprotease (Roche), 0.5mM Na3VO4, 20mM beta-Glycerophosphate, 0.1mM DTT, 1U/ul RNasin (Promega). Cells were removed from the plate by scraping, and pelleted by centrifugation at 1600 rpm for 5 minutes at 4°C. Cell pellets were resuspended in 400 µL of lysis buffer (SW buffer containing 10% glycerol and 0.1% NP-40) and gently pipetted up and down 15 times using a P1000 tip and centrifuged at 2500 rpm for 5 minutes at 4°C to pellet the nuclei. The supernatant was considered as the cytosolic fraction. Nuclear pellets were resuspended in 400 µL of RB buffer containing 9 mM EDTA, 0.2 mM EGTA, 0.1% Triton X-100, supplemented before use with 1x antiprotease (Roche), 0.5mM Na3VO4, 20mM beta-Glycerophosphate, 1mM DTT, 1U/ul RNAsin (Promega) incubated for 15 minutes while rotating at 4°C, and centrifuged at 3000g for 5 minutes to pellet chromatin. The supernatant was considered as the nucleosolic fraction. Chromatin pellet were resuspended in 400 µL of RB buffer supplemented with 1% SDS and solubilized by sonication using a water bath sonicator (Diagenode) at high amplitude (6 cycles of 20 seconds). In parallel, total cell extracts were prepared in RIPA buffer: 150 mM NaCl, 50 mM Tris pH 8, 0.25% sodium deoxycholate, 0.1% SDS, 1% NP-40, freshly supplemented with 0.5 mM DTT and 1× protease inhibitor, RNAsin then incubated for 10 minutes on ice.

### Co-immunoprecipitation assays

Approximately 10⁶ cells from a confluent 100 mm plate were used to prepare nuclei according to the cell fractionation protocol described above. Nuclear pellets were resuspended in 0.5 ml of RIP buffer: 25mM TRIS pH 7.4, 150mM KCl, 2.5% glycerol, 0.3% NP-40 freshly supplemented with 1x antiprotease, 0.5 mM Na3VO4, 20mM beta-glycerophosphate, 0.1 mM DTT, 1U/ul RNAsin, and treated with 7U/mL TURBO DNase. The suspension was incubated at 37°C for 5 minutes. Samples were sheared by three cycles of sonication (10 seconds ON, 10 seconds OFF) using a Bioruptor (Diagenode), followed by a second incubation at 37°C for 5 minutes. The reaction was stopped by addition of 10 µl /mL of 0.5M EDTA per 0.5 ml. Samples were then centrifuged at 12000g for 10 minutes at 4°C. The resulting supernatant, referred to as “RIP extract”, was used for subsequent immunoprecipitation (IP) experiments, and a portion of the RIP extract was reserved as input samples.

For each IP, 150 µl of RIP extract was combined with 3 µg of antibodies and 350 µl of RIP buffer. The mixture was incubated for 1h30 at 4°C on a rotating wheel. Saturated magnetic beads (Dynabeads M-280 sheep anti-rabbit IgG or Dynabeads Protein G, Invitrogen) were prepared by blocking 80 µl of beads per IP in PBS supplemented with 0.1% BSA, 0.5% NP40, and 0.1% polyvinylpyrrolidone (PVP) for 30 min at 4°C, followed by two washes with Wash V buffer: 0.1% NP-40, 150mM NaCl, 20 mM TRIS.Cl pH8, 1mM EDTA. The saturated beads were resuspended in 300 µl of Wash V buffer per IP, and added to the IP mixture. After 45 minutes of incubation, beads were then washed twice with RIP buffer, twice with Wash V buffer, and once with TE/NP40 buffer: 10 mM TRIS pH7.4, 1 mM EDTA, 0.01 % NP-40. Beads were eluted in Laemmli sample buffer. Both eluates and input samples were further processed for Western blot analysis.

### Cell fractionation for RNA preparation and sequencing (RNAseq)

For total RNA extraction, cells were trypsinized, and the resulting pellet was washed with PBS, and pelleted by centrifugation at 1800 rpm for 4 minutes at 4°C. The cell pellet was resuspended in 1600 µL of Buffer D: 4 M guanidinium thiocyanate, 25 mM sodium citrate, pH 7.0, 0.5% (wt/vol) N-laurosylsarcosine and 0.1 M 2-mercaptoethanol ^60^, homogenized thoroughly for total cell extracts, from which the total RNA was extracted.

For chromatin-associated and cytosolic RNA extraction, cells were resuspended, as described above for cell fractionation, in 125 µL of ice-cold SW buffer and incubated on ice for 8 minutes. After swelling, 125 µL of lysis buffer was added, and the suspension was gently pipetted 15 times before centrifugation at 3500 rpm for 4 minutes at 4°C. The nuclei pellet was resuspended in 250 µL of Buffer RB and incubated on ice for 15 minutes, followed by centrifugation at 6000 rpm for 4 minutes at 4°C, separating the chromatin and nucleosol. Both the nucleosol and chromatin fractions were homogenized in Buffer D for RNA extraction.

RNA was extracted by addition to all fractions of 250 mM sodium acetate, 1 volume of acidic phenol, and 0.2 volumes of chloroform. The samples were incubated on ice for 10 minutes, followed by centrifugation at 14,000 g for 20 minutes at 4°C. The supernatant was collected and mixed with an equal volume of chloroform, then centrifuged at 14,000 g for 5 minutes at 4°C. The resulting supernatant was combined with an equal volume of isopropanol, vortexed thoroughly, and incubated overnight at -20°C for RNA precipitation. The samples were centrifuged at 14,000 g for 20 minutes at 4°C, and the resulting pellet was washed with 75% ethanol, and resuspended in 100 µL of Buffer D (diluted 1:10), supplemented with 0.5% Triton X-100, 0.4 M NaCl, and 1 µL of proteinase K. The mixture was incubated at 50°C with shaking at 850 rpm for 3 hours. Subsequently, 250 mM sodium acetate, 300 µL of Buffer D, and 400 µL of acidic phenol/chloroform were added to each fraction. The samples were centrifuged at 14,000 g for 15 minutes at 4°C, and the supernatant was collected. An equal volume of chloroform was added to the supernatant, followed by centrifugation at 14,000 g for 20 minutes at 4°C. The final supernatant was supplemented with Glycogen (2 µg/mL) as carrier and 1 volume of isopropanol, and the samples were precipitated overnight at -20°C. The pellet was then washed with 75% ethanol and resuspended in 50 µL of RNase-free water. The samples were incubated at 45°C with shaking at 800 rpm for 10 minutes. 40 µL was subjected to DNase treatment for 50 minutes at 37°C. RNA integrity was verified by agarose gel electrophoresis. On all RNA fractions, total RNA library preparation and sequencing were performed by Novogene (UK) Co., Ltd as part of their lncRNA-seq service, which included directional library preparation with rRNA removal and sequencing on the Illumina NovaSeq platform using paired-end 150 bp (PE150) reads.

### Chromatin immunopreciptation and sequencing (ChIPseq)

BMEL cells were crosslinked following a modification of the protocol described in ref. ^61^. Briefly cells were incubated in freshly prepared 2 mM DSG (Merck) in PBS for 50 minutes at room temperature with occasional shaking. After three washes in PBS, cells were incubated for 10 minutes in 1% formaldehyde (Merck) in PBS. The crosslinking reaction was quenched with 125 mM glycine for 5 minutes at room temperature. Cells were washed with PBS, harvested, and pelleted by centrifugation at 3000 g for 3 minutes. Cells were subsequently resuspended in 1.4 mL of freshly prepared swelling buffer (25mM Hepes pH 7.95, 10mM KCl, 10mM EDTA) supplemented with protease inhibitors (Roche) and 0.5% NP-40. The suspension was incubated on ice for 30 minutes, then pipetted up and down 50 times using a P1000 pipette tip. The cells were centrifuged at 6000g for 2 minutes and resuspended in 300 µL of TSE150 buffer with 0.3 % SDS (0.3% SDS, 1% Triton, 2mM EDTA, 20mM Tris-HCl pH8, 150mM NaCl) freshly supplemented with protease inhibitors. Chromatin was sonicated in 1.5 mL tubes for a total of 15 minutes using cycles of 30 seconds ON and 30 seconds OFF, with the samples maintained in ice-cold water. Following sonication, the samples were centrifuged at 14,000g for 30 minutes at 18°C. An aliquot of supernatant was used to quantify chromatin and check DNA fragment size (typically around 300 bp).

Chromatin immunoprecipitation (ChIP) was performed using 70 µg of chromatin and 4 µg of the following antibodies : anti-MTR4/SKIV2L2 (Bethyl A300-615A), anti-ZC3H18 (Atlas HPA 040847), anti-EXOSC10 (Bethyl A303-987A), anti-CTCF (Diagenode A2354-0023P), anti-P-RNAPII (a mix of antibodies against phospho-Ser5 and phospho-Ser2, Abcam Ab5095 and Ab5408, respectively), and anti-murine IgG (Merck). Immunoprecipitation was carried out overnight on a rotating wheel at 4°C. Subsequently, 90 µL of Protein G beads (Dynabeads, Invitrogen), blocked overnight at 4°C in blocking buffer containing 5% Buffer C (0.5% SDS, 10 mM Tris-Cl pH 8, 1 mM EDTA, 0.5 mM EGTA), 20% Buffer D 1.2× (1.2% Triton, 0.12% NaDoc, 180 mM NaCl, 12 mM Tris-Cl pH 8, 1.2 mM EDTA, 0.6 mM EGTA), 72% Wash Buffer V, 1.5% BSA (Merck), and 1.5% Polyvinylpyrrolidone (Merck), were added to the IP samples and incubated for 1h on a rotating wheel at 4°C. Ten percent of the chromatin was set aside as input. Following immunoprecipitation, the beads were pelleted and subjected to sequential washes, each performed for 3 minutes on a rotating wheel at room temperature using 1 mL of the following buffers: Three washes in TSE150 buffer with 0.1 % SDS (0.1% SDS, 1% Triton, 2mM EDTA, 20mM Tris-HCl pH8, 150mM NaCl), then one wash in TSE500 buffer (TSE150 buffer with 0.1% SDS and 500 mM NaCl), then one wash in buffer (10 mM Tris-HCl pH 8, 0.25 M LiCl, 0.5% NP-40, 0.5% sodium deoxycholate, 1 mM EDTA), then two washes in TE buffer (10 mM Tris-HCl pH 8, 1 mM EDTA). Elution was performed in 300 µL of elution buffer (1% SDS, 10 mM EDTA, 50 mM Tris-HCl pH 8) by vortexing vigorously, followed by incubation at 65°C for 15 minutes. A second elution was performed using 100 µL of elution buffer. Both eluates were collected, and the beads were rinsed in 100 µL of TE buffer with 1% SDS, subsequently added to the pool of eluates. Both the immunoprecipitated and input chromatin samples were incubated overnight at 65°C to reverse crosslink, followed by proteinase K and RNase A treatments. DNA was isolated using a NucleoSpin PCR purification kit (Macherey Nagel), and concentrations were measured using a Qubit fluorometer (Invitrogen). Purified ChIP samples were processed for Illumina paired-end (PE40) sequencing performed at the Genom’IC core facility (Paris, France).

### Western Blotting analysis

Protein samples were denatured by the addition of Laemmli sample buffer (Bio-Rad) with reducing agent at 100°C for 10 minutes and separated by SDS-PAGE (Bio-Rad, Criterion XT), then transferred to a nitrocellulose membrane. Transferred proteins were visualized by Ponceau S staining, and the membranes were incubated overnight at 4°C in PBS-T (PBS, 0.1% Tween 20), containing 5% (w/v) non-fat dry milk. The membranes were then incubated for 1h30 at room temperature with primary antibodies at the appropriate dilution in PBS-T (as described in the Western blot antibodies section). After three 5-minute washes in PBS-T, the membranes were incubated for 1h30 with secondary antibodies (Starbright Blue 700 anti-rabbit/mouse IgG (Biorad), or rabbit true blot (Rockland), all diluted 1/4000), followed by three additional 5-minute washes. Protein detection was performed either by fluorescence at 700 nm or by chemiluminescence using the Pierce™ ECL Plus Western Blotting Substrate (Thermo Fisher Scientific) and visualized with a ChemiDoc MP imaging system (Bio-Rad).

### Western blot antibodies

| Antibodies | Dilution | References |
| --- | --- | --- |
| Anti-EXOSC3 | 1/1000 | Proteintech 15062-I-AP |
| Anti-EXOSC9 | 1 /1000 | Abcam ab156686 |
| Anti-EXOSC10 | 1/1000 | Bethyl A303-987A |
| Anti-MTR4/SKIV2L2 | 1 /1000 | Bethyl A300-615A |
| Anti-ZC3H18 | 1 /2000 | Bethyl A304-682A |
| Anti-CTCF | 1/1000 | Diagenode A2354-0023P |
| Anti-HP1α | 1 /1000 | Euromedex 2G9 |
| Anti-HP1β | 1/2000 | Euromedex 1A9 |
| Anti-HP1γ | 1/3000 | Euromedex 1G6 |
| Anti-H3K9me3 | 1/2000 | Abcam ab8898 |

### ATACseq

Chromatin accessibility assays were performed on each of the six Het and TKO BMEL clones according to the protocol described in ref. ^62^, with an ATACseq kit (ActiveMotif, 53150). Paired-end (PE40) sequencing of the dual-index ATACseq libraries was performed by Novogene (UK) Co., Ltd.

### CUT&Tag

CUT&Tag assays were performed on two independent biological replicates of Het and TKO BMEL cells according to the protocol by ^63^, but with the NE1 buffer (https://www.protocols.io/view/bench-top-cut-amp-tag-kqdg34qdpl25/v3) to permeabilize cells, and with the same anti-HP1α, -HP1β, -HP1γ, -H3K9me3 as used in Western blot analyses, and with anti-H3K27me3 (Cell Signaling Technol. 9733). Libraries were obtained with 15-16 PCR amplification cycles such that at least a faint ladder of nucleosome-sized bands were visible using an Agilent Tape Station High-Sensitivity D1000 Screen Tape representing DNA concentrations of at least 200 pg/µl for anti-HP1 CUT&Tag reactions with Het cells, and at least 10-fold lower concentrations for the same reactions using TKO cells. 40 nt paired-end sequencing was performed at the I2BC sequencing platform. Reads were aligned on the mm10 reference genome with bowtie2 (parameters: --end-to-end --very-sensitive --no-mixed --no-discordant -k 1 -X 1000 --phred33 -I 25 -p 24 -x mm10). Bigwig files were generated from two merged HP1 CUT&Tag replicates, and they were normalized using CPM.

### Bioinformatics analyses

RNAseq data analysis : Mapping was carried out with STAR (v2.6.0b) (parameters: – outFilterMismatchNmax 1 – outSAMmultNmax 1 –outMultimapperOrder Random – outFilterMultimapNmax 30). The reference genome was GRCm38 Mus musculus primary assembly from Ensembl. The SAM files were converted to BAM files and sorted by coordinate using samtools (v1.8). Gene expression analysis was performed with the DESeq2 (v1.18.1) package. P-values from the differential gene expression test were adjusted for multiple testing according to the Benjamini and Hochberg procedure.

ChIPseq data analysis : Raw ChIP-seq data in Fastq format were subjected to quality control using FastQC (v0.11.9). ChIP-seq reads were mapped to mm10 using bowtie2 (v2.3.4) (parameters: -N 0 -k 1 –very-sensitive-local). We then selected reads with a MAPQ equal or higher than 30 corresponding to uniquely mapped reads for further analysis. Peak calling was performed using MACS2 (v.2.1.1) (parameters: -p 0.05).

ATACseq data analysis : For chromatin accessibility profiling, Bowtie2 was used with the parameters --very-sensitive -X 2000 to accommodate the broad range of ATAC-seq fragment sizes, and --maxins 500 --no-discordant --no-mixed -k 1 to eliminate multi-mapping reads. Aligned BAM files were sorted and indexed using SAMtools (v1.8). ATAC-seq peaks were called using MACS2 (v2.2.9.1) with the parameters --nomodel, --shift -100, and --extsize 200, which are recommended for ATAC-seq to account for the Tn5 transposase cutting pattern. Peaks were identified using a q-value threshold of 0.01 (-q 0.01), and signal tracks were generated in bedGraph format with -B --SPMR. The genome size was set to mouse (-g mm), and input files were sorted BAM files in BAM format (-f BAM).

For all high throughput experiments, Bigwig files were generated from BAM files with bamCoverage (parameter: –normalizeUsing CPM) from Deeptools (v3.1.3)

Pathway analyses with gene names were performed with Enrichr (https://maayanlab.cloud/Enrichr/). The FDR (adjusted p-value, denoted p-adj in Figures) of each path entry served as the significance threshold.

### Definition of genomic regions and categories

LTR, SINE, LINE, and MMSAT regions were defined by the RepClass field in the Repeat masker database in the UCSC table browser (https://genome.ucsc.edu/). The enhD Distal enhancers were defined as the dELS/enhD category in the Encyclopedia of DNA Elements (ENCODE) candidate cis-regulatory element (cCRE) database in the UCSC table browser. Extragenic repeats and enhD elements were defined relative to genes in the Ensembl Grc38 database (https://www.ensembl.org/). Promoter associated transcripts (PROMPT) regions were defined as 2kb upstream of transcription start sites (TSS) of genes. Exons were defined according to the Ensembl GRCm38 database. The list of 23808 Genes was defined according to the Ensembl Grc38 database, after removing all the entries with names beginning with “Mir”, “Rik”, and “Gm”, for the sake of clarity of the average density profiles.

The EnhD (Up, Down, Unchanged, and Closed) categories were created according to their activity as defined by ATACseq read counts obtained by featurecounts on distal enhancer elements (enhD) in ENCODE database. EnhD categories are as follows :

enhD Up (ATACseq reads>10 p-val.<0.05 FC TKO vs. Het >=4),
enhD Down (ATACseq reads >10, p-val.<0.05 FC TKO vs. Het <= 0.25),
enhD Unchanged (ATACseq reads >10, 0.9<FC<1.1),
enhD Closed (ATACseq reads <2).

### RNA Read counts on genomic regions

Quantification of uniquely mapped reads from the transcriptome was carried out with featureCounts (v1.6.1) from the Subread suite ^64^, with the following parameters: -p -B -C -s 1 -- minOverlap 25. FeatureCounts with these parameters counted only uniquely mapped reads with both paired-ends aligned (-p -B).

Chromatin and cytoplasmic stranded RNA reads were counted on extragenic repeats only in order to evaluate self-expressed repeats and avoid counting reads on gene-embedded repeats which would reflect expression of the surrounding gene and not the repeat itself. Read counts were the results of stranded reads counted in regions substracted by stranded reads counted in the 200nt region upstream of repeats adjusted for their size.

Transcriptome reads on PROMPT were counted antisense, corresponding to upstream-antisense RNA (uaRNA), relative to the orientation of their adjacent gene substracted by antisense read counts in its first exon adjusted for their respective size. Transcriptome read counts on enhD were substracted by read counts in the highest of the two adjacent 200nt region upstream or downstream of each enhD adjusted for their respective size. All genomic regions counted for RNA levels were considered expressed when two or more reads (or ten or more reads on exons) were counted in average of the triplicate samples, in either Het or TKO or both, after substracting reads from size-normalized adjacent regions (illustrated in Fig. 2B and 2E). In order to avoid excluding the elements expressed only in Het or in TKO, a pseudo-count of 0.1 was added to the mean RNA read counts before transformation into Log2 (illustrated in Fig. 2C, 2D, 2F and 2G).

### Data visualization

BigWig files of samples or merged replicates were visualized with the Integrative Genomics Viewer (IGV version 2.8.10). Heatmaps and profiles were generated with the computeMatrix, plotProfile and plotHeatmap tools of the Deeptools package (v3.1.3). The patient survival curves in Supplementary Fig. 6B are based upon data generated by the TCGA Research Network: https://www.cancer.gov/tcga, obtained from the Human Protein Atlas database (proteinatlas.org).

### Statistical analyses

The statistical tests performed are described in Fig. legends or in the Methods section. Significance level was evaluated by two-tailed Student’s t-test as indicated in Fig. legends. The exact (n) number of samples is indicated in each figure. No statistical method was used to predetermine sample size. No data were excluded from the analyses.

### Data availability

All RNA-seq, ChIP-seq, ATAC-seq, and CUT&Tag datasets generated during this study are available in the EBI BioStudies database (http://www.ebi.ac.uk/biostudies) under accession numbers: E-MTAB-15194, E-MTAB-15196, E-MTAB-15197, and E-MTAB-15189, respectively.

## Acknowledgments

The authors wish to thank Enzo Becherel for its technical help. This work has been supported by PhD fellowships from La Ligue Nationale Contre le Cancer (to HS) and by grants from Aviesan ITMO Cancer (Plan Cancer, AAP ARN non-codants) and Institut National du Cancer (INCa, PLBIO-2020-107). We acknowledge the sequencing and bioinformatics expertise of the I2BC High-throughput sequencing facility, supported by France Génomique (funded by the French National Program “Investissement d’Avenir” ANR-10-INBS-09).

## Author Contributions

Experimental investigation: H.S., L.S., K.G., E.B., F.C., C.Ma., C.R.

Bioinformatic analysis: H.S., M.C., C.Ma., C.Mu., C.R.

Funding acquisition: C.Mu.

Writing original draft: H.S., C.Mu., C.R.

Manuscript review and editing: F.C., C.Ma., C.Mu., C.R.

All authors read and approved the final version of the manuscript.

## Competing Interests

The authors declare no competing interests.

## Material availability

The Het and TKO BMEL cell lines are readily available from the authors upon request.

---

**Supplementary Fig. 1:**
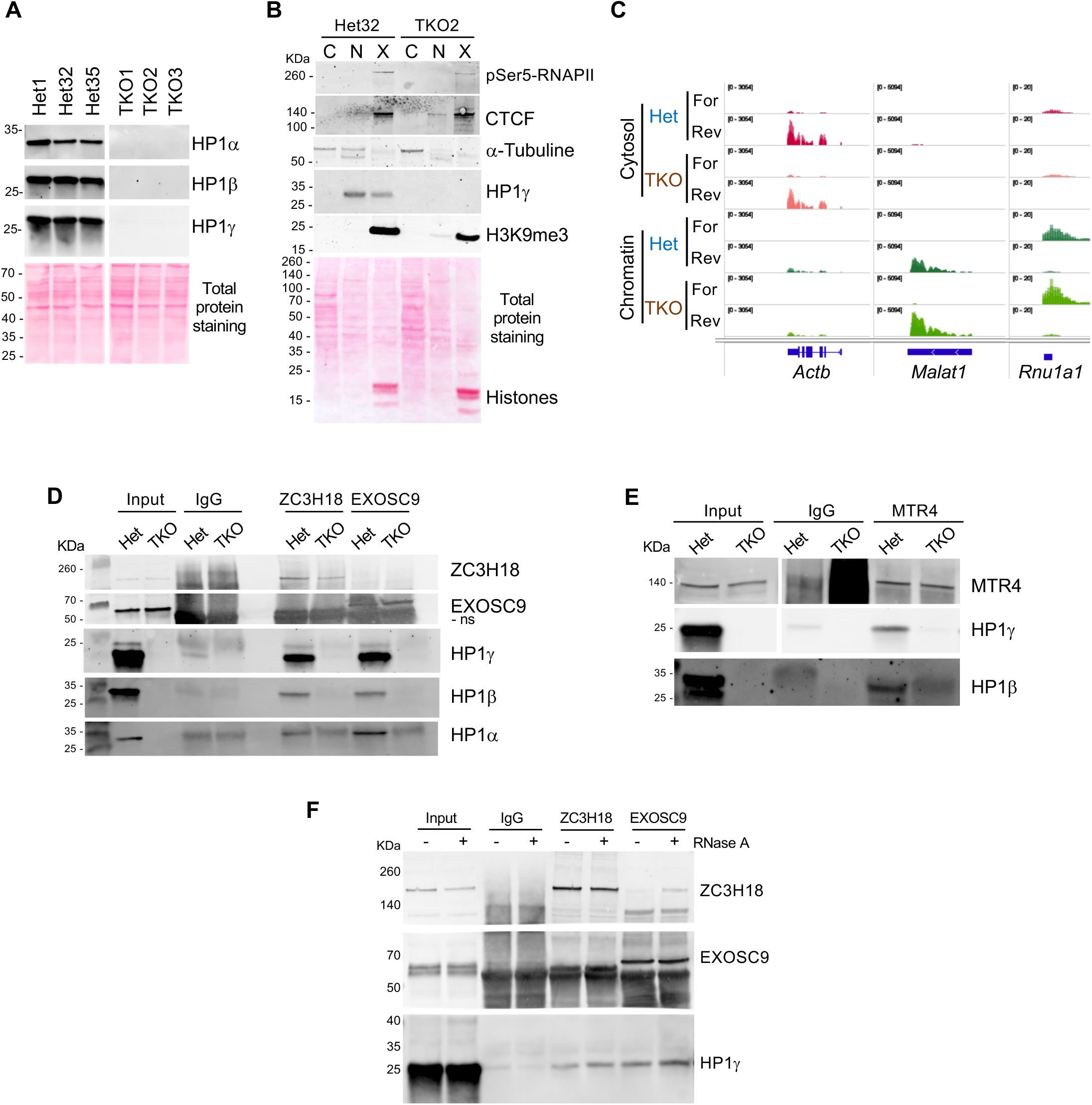
Cell fractionation and co-immunoprecipitation assays. **A.** Levels of HP1 proteins in all BMEL clones by Western blot analysis. **B.** Distribution of indicated proteins in the subcellular fractions (C, N, X, as in Fig. 1B) in two representative Het and TKO clones by Western blot analysis, and total proteins by Ponceau S staining. **C**. Genome views as in Figure 2A of the distribution of RNA density profiles between cytosolic and chromatin fractions for Actb mRNA and the nuclear Malat1 RNA or U-snRNA. Distribution reflects the expected subcellular localizations of the depicted RNA species. **D, E.** Co-immunoprecipitation assays of HP1a, HP1b, and HP1g by MTR4, ZC3H18, or EXOSC9, compared to IgG. **F.** RNA-independent co-immunoprecipitation of HP1g by ZC3H18 and EXOSC9. HP1g signal is not lost upon RNase A treatment.

**Supplementary Fig. 3:**
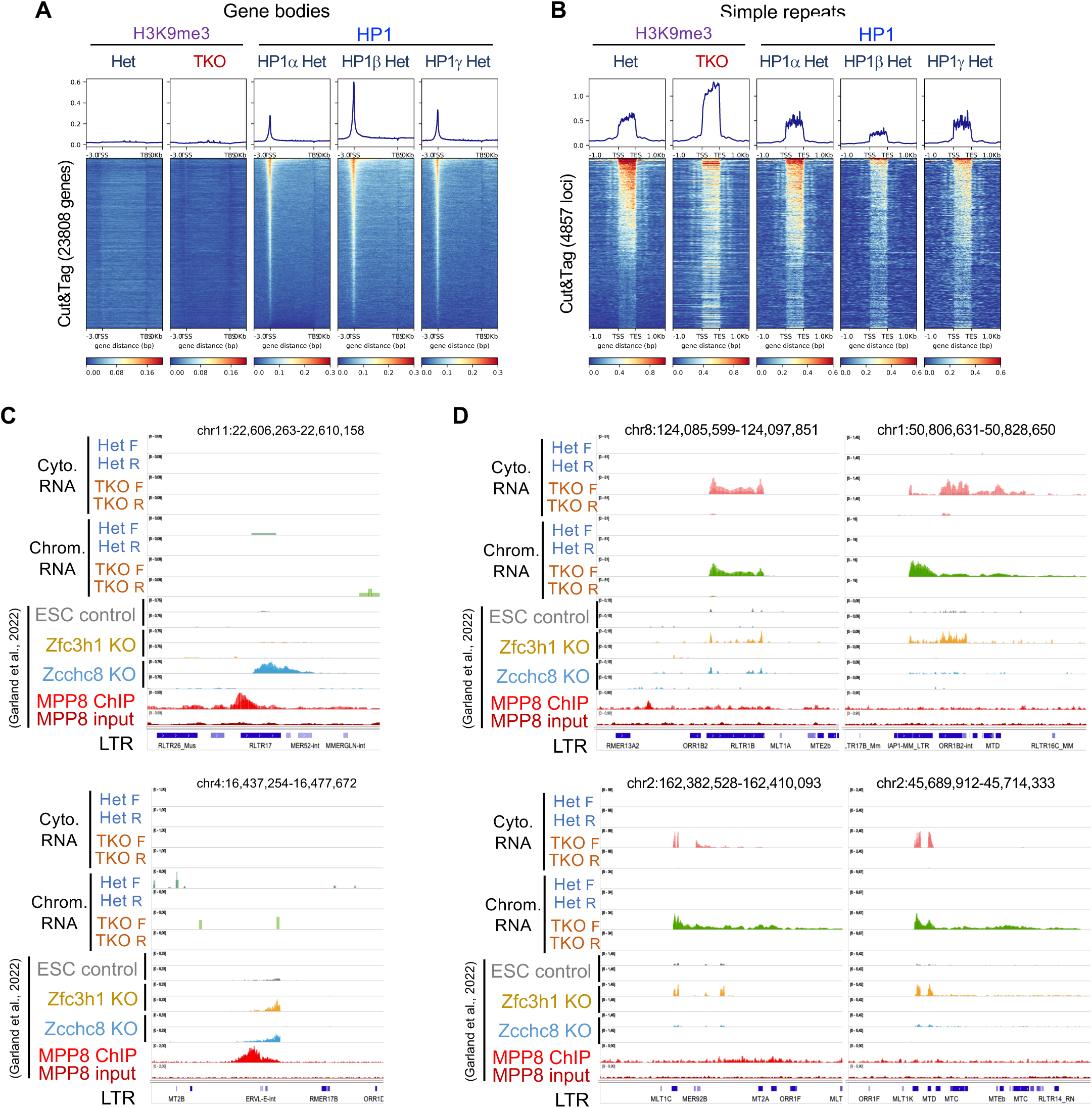
HP1/RNA Exosome loci and H3K9me3 and MPP8 components of the HUSH complex. **A,B.** Heatmaps of densities of H3K9me3 or HP1 isoforms by Cut&Tag in Het or TKO as indicated, on gene bodies (A) as in figures 3D-F, or on a list of simple repeats (B) from Repeat Masker database. **C, D.** Genome views of transcriptomes in Het or TKO (examples depicted in figure 2H) compared with transcriptomes of mESC invalidated for the expression of *Zcchc8* KO (sky blue profiles) or *Zfc3h1* KO (yellow profiles) and control mESC (grey profiles), together with MPP8 chIPseq (red) and input (dark red) profiles (GSE178550; Garland et al., 2022). **C.** Genome views of two examples of MPP8 ChIPseq peaks with a coincidental RNA profile in mESC KO. No upregulated RNA was detectable at these loci in Het or TKO. **D.** Genome views of upregulated RNA (green and red) in TKO at LTR loci, with a coincidental upregulated RNA in mESC KO (yellow and sky blue). No MPP8 ChIPseq peak was detectable at these loci.

**Supplementary Fig. 5:**
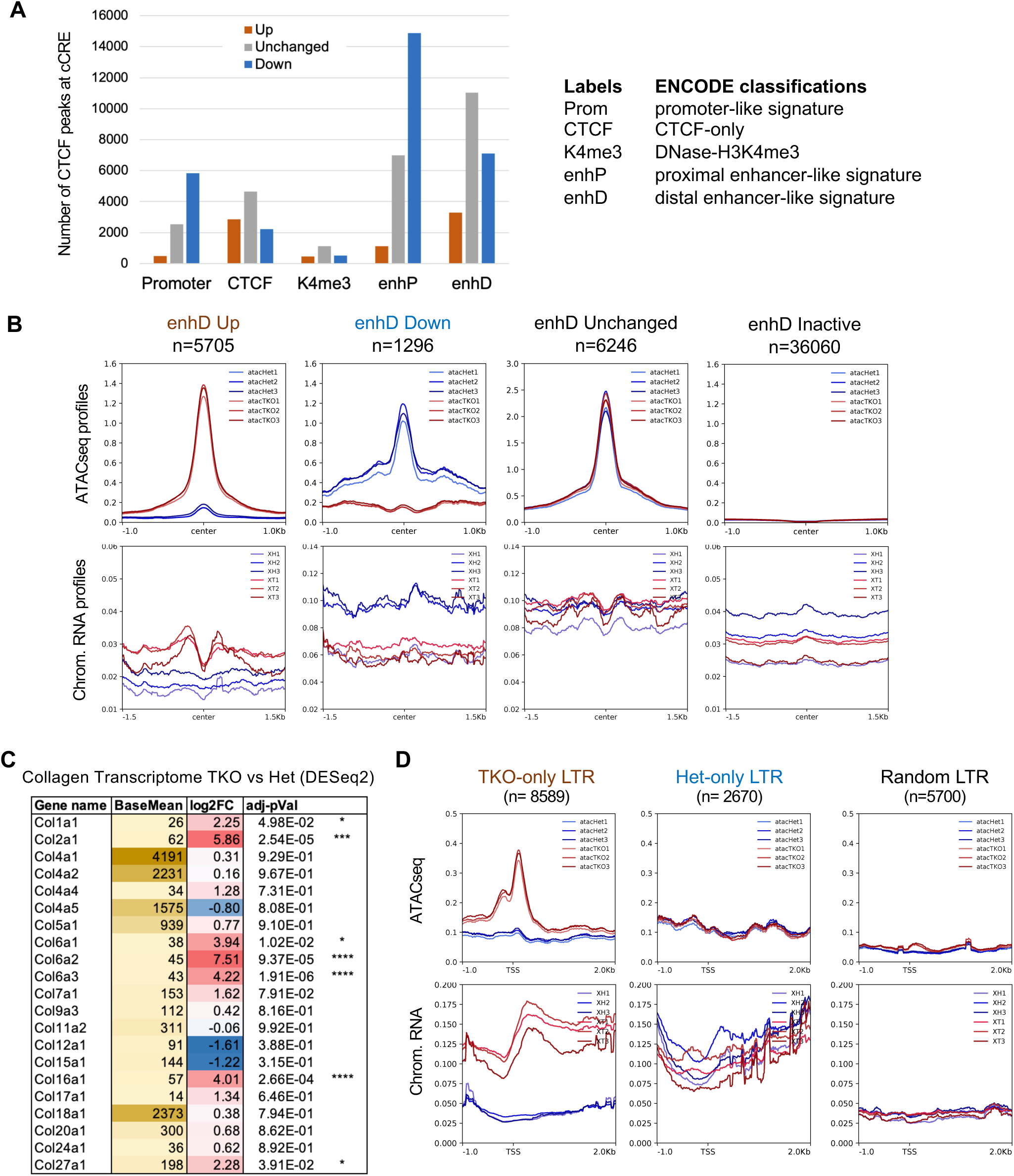
ATACseq and chromatin-associated RNA at enhancers and LTRs. **A.** Number of CTCF ChIPseq peaks located at the cCRE regulatory elements labelled according to the ENCODE cCRE classifications (right panel). Bars represent CTCF peaks that were up- (brown) or down-regulated (blue) by more than 2 fold in TKO versus Het, or changed by less than 20 percent (Unchanged, grey). **B.** Average density profiles on enhD elements. Top panels, ATACseq profiles, and bottom panels, chromatin-associated RNA density profiles in triplicate Het (blue profiles) and TKO (red profiles) samples on enhD elements in the cCRE ENCODE database. EnhD were categorized based on their chromatin accessibility by ATACseq between Het and TKO. Categories are as follows: Down, with decreased accessibility in TKO cells; Unchanged, active in both conditions; Up, with increased accessibility in TKO cells; inactive, with no ATACseq density, respectively. **C.** List of Collagen genes dysregulated in Het versus TKO by DESeq2 analysis. Base mean is their mean expression levels in both Het and TKO, log2FC is the fold change in TKO versus Het, with adjusted p-values. **D.** Average density profiles on LTR retroelements. Top panels, ATACseq profiles, and bottom panels, chromatin-associated RNA density profiles in triplicate Het (blue profiles) and TKO (red profiles) samples. TKO-only LTRs and Het-only LTRs correspond to the categories of extragenic expressed LTRs depicted in Fig. 2B. Random LTR is a subset of 5,700 LTR elements in the Repat Masker database chosen randomly.

**Supplementary Fig. 6:**
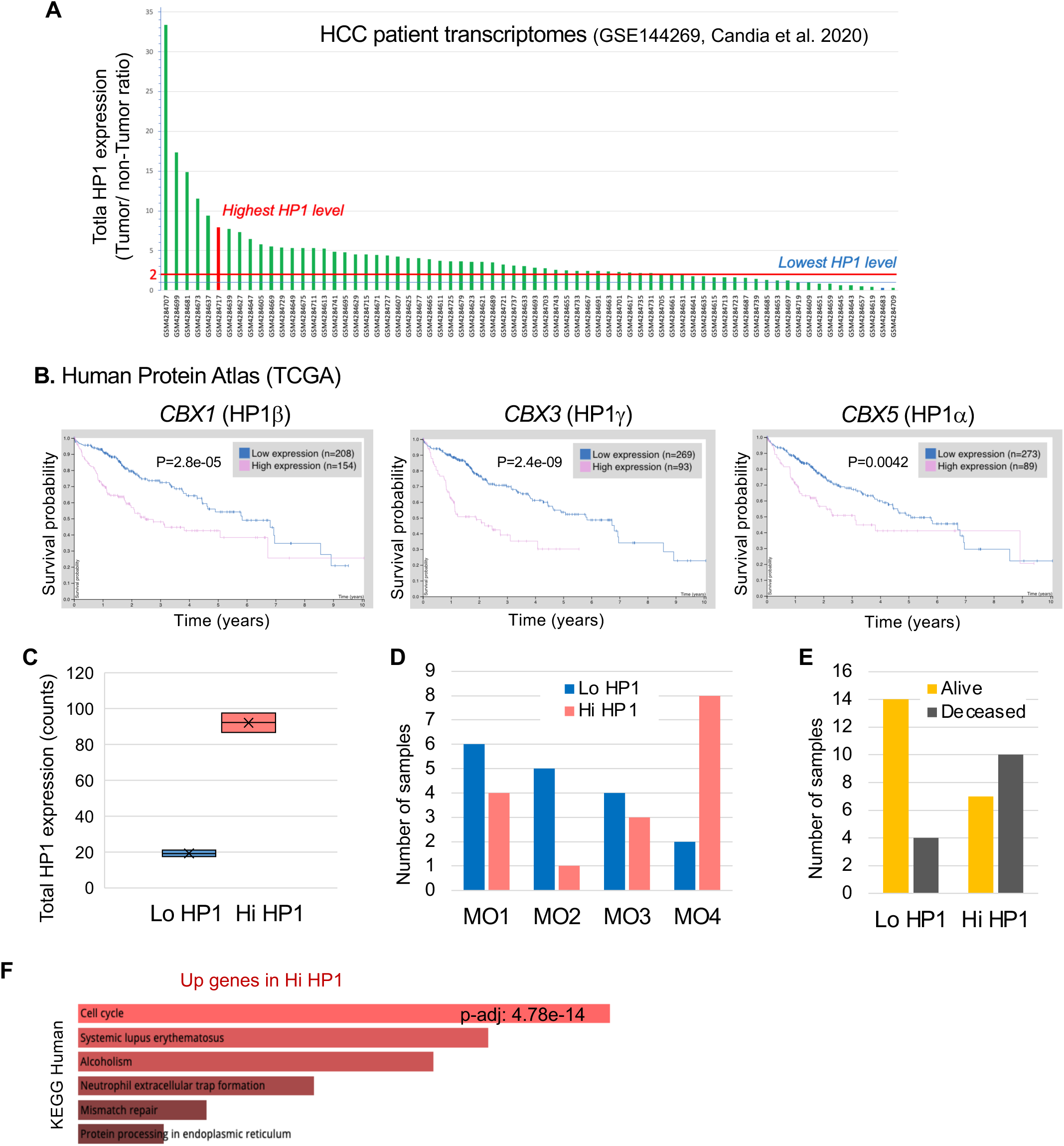
Metadata associated with the analysis of HCC patient samples. **A,**Histogram of total HP1 expression ratio in tumors relative to adjacent non-tumor samples of 76 HCC patient transcriptomes (GSE144269, ref. 47) **B.** Kaplan-Meyer plots of liver cancer patient survival probabilities associated with *CBX1* (HP1b), *CBX3* (HP1g), and *CBX5* (HP1a) from the. Survival curves for low expression (blue) are compared to high expression (pink) of the indicated genes. P scores indicate log-rank P values of correlation between expression levels and patient survival. **C.** Total HP1 expression levels in the two groups of high HP1 (Hi HP1) and Low HP1 (Lo HP1) patients defined by the first and third total HP1 expression quartiles, respectively, related to Fig. 6A. **D, E,** Histograms of the number of Hi HP1 and Lo HP1 patients clustered according to four Molecular subclasses (MO) sorted by increasing severity of the disease from MO1 to MO4 (D), or showing survival criteria (E), as defined in ref. 47. **F.** Histograms of the most significant KEGG (Kyoto Encyclopedia of Genes and Genomes) pathways corresponding to upregulated genes in Hi HP1, sorted by adjusted p-values.

## Notes

### Competing Interest Statement

The authors have declared no competing interest.

### Summary of Updates

This version of the manuscript has been revised to update manuscript format.

